# Evolution of immune escape mechanisms in the progression from preinvasive to invasive human lung adenocarcinoma

**DOI:** 10.1101/2020.04.17.046540

**Authors:** Nasser K. Altorki, Alain C. Borczuk, Vivek Mittal, Olivier Elemento, Timothy E. McGraw

**Author notes:** co-corresponding authors: Nasser K Altorki, 525 East 68^th^ Street Room M404, New York, NY 10065, NKA, Timothy E McGraw, 1300 York Ave, Room W305, New York, NY 10065, TEM.

## Abstract

The tumor microenvironment (TME) of lung adenocarcinoma (LUAD) precursor lesions has not been described. We interrogated by multiplex immunofluorescence the TME of preinvasive and invasive Stage 1A LUADs selected by computer tomography (CT) scan-density. Pure non-solid (p-NS) CT density nodules are preinvasive/minimally invasive, whereas solid CT density nodules are frankly invasive cancers. Our data reveal an intensely immune-suppressive immune TME in p-NS tumors characterized by an increase in Treg cells and a decrease in cytotoxic T cells relative to normal lung. The TME of the solid tumor group, more advanced lesions than the p-NS yet still early in disease development, were increasingly more immune-suppressive. Provocatively, there was a further increase in both Treg cells and cytotoxic T cells, establishing a nascent albeit ineffective anti-tumor immune response in transition from preinvasive p-NS to invasive solid tumors. Regulatory T cells play a dominant role throughout progression, while additional immune evasive mechanisms are employed at different stages of disease progression, including T cell exclusion from cancer cell nests early and activation of immune checkpoints later. Our study establishes that different immune-targeted strategies are required to intercept disease progression at these two distinct early points of lung cancer development.

**Statement of Significance:** Using multiplexed IF, we compared the cellular composition and activation state of the tumor immune microenvironment between pre/minimally invasive and frankly invasive adenocarcinoma. We found a progressive increase in immunosuppressive mechanisms in association with disease progression suggesting that Interception strategies should be specifically tailored based on underlying immune escape mechanisms

## INTRODUCTION

The widespread use of computerized chest tomography (CT), particularly for lung cancer screening purposes, has led to the detection of ground-glass or non-solid nodules that are not visible on plain chest radiography. Non-solid nodules appear on CT scanning as hazy opacities that do not obscure the underlying lung parenchyma or vasculature. In most instances, non-solid nodules less than 5 mm in size represent focal proliferative lesions known as atypical alveolar hyperplasia that are considered the earliest progenitor lesions of invasive adenocarcinoma of the lung. The majority of non-solid nodules, particularly those ≤ 3cm in size, typically harbor adenocarcinoma *in-situ* (AIS) or minimally invasive adenocarcinoma (MIA) where the invasive component is ≤ 5 mm or less. Interestingly, 10-20% contain adenocarcinoma with an invasive component exceeding 5 mm suggestive of a more aggressive phenotype. Regardless, the precise biological behavior of these non-solid nodules remains unclear as many remain unchanged in size and appearance for many years. However, approximately 20-40% of these nodules grow or develop areas of increased CT density (a manifestation of more invasive malignancy) within a four-year window (1). Why some of these nodules retain an indolent behavior while others progress to invasive malignancy is unclear. This uncertainty commits patients to frequent, repeat imaging, and in some cases potentially harmful biopsies.

The critical role of the tumor microenvironment (TME) in tumorigenesis is well-recognized (2-5). The TME is predominantly composed of immune cells, fibroblasts and vasculature. There are a number of non-mutually exclusive hypotheses for how the TME contributes to tumorigenesis, one of which is “immunoediting” or “immune surveillance”, a cancer cell-extrinsic mechanism that is engaged once cell-intrinsic killing mechanisms (e.g., apoptosis) have failed to eliminate individual transformed cells (6-8). The proposed temporal phases of cancer immune surveillance describing the homeostasis between cancer cell killing and survival include: elimination, equilibrium and escape (9). In the elimination phase, innate and adaptive immunity eliminate cancer cells before a tumor can emerge. In the equilibrium phase, immune cell killing of cancer cells is balanced by cancer cell proliferation, such that tumor growth is maintained in check, but the cancer cells are not fully eliminated. The escape phase is one of tumor growth due to cancer cells evading immune cell-mediated killing. Entry into the escape phase portends disease progression.

There are multiple mechanisms that may contribute to tumor progression, including the development of an overall immunosuppressive state in the TME and/or cancer cell driven escape mechanisms such as antigen loss or aberrations that impede antigen presentation such as downregulation of MHC class I or II molecules. Another mechanism invoked for immune evasion is T cell exclusion from cancer cell nests attributed to organized fibroblastic proliferation leading to physical and/or chemical barriers that restrict access of T cells to the cancer cell nests. Any or all of the above mechanisms of immune escape may contribute to tumor progression in a temporal and context dependent manner.

We posited that progression of lung cancer from preinvasive to invasive adenocarcinoma develops as a result of immune escape mediated by a shift towards a more immune-suppressive microenvironment. To test this hypothesis, we used multiplex immunofluorescence to compare the cellular compositions and activation states of the TMEs of radiographically pure non-solid nodules (preinvasive or minimally invasive lung cancer) to the TMEs of radiographically solid (invasive lung cancer) nodules. Given the importance of CT appearance in driving clinical decisions in management of lung nodules, we classified the groups based on their radiographic appearance (10). Comparison of these two tumor groups captures changes in the TME correlated with progression from preinvasive pure non-solid (p-NS) to invasive solid tumors. The characteristics of the TMEs of both p-NS and solid tumors are progressively more immune suppressive than adjacent normal lung. Our findings demonstrate a diversity of immune escape mechanisms at different phases during early development of human lung adenocarcinoma (LUAD).

## Methods

### Samples and Tissue microarray

The samples for this IRB approved study were collected from the Weill Cornell lung nodule cohort, a retrospective collection of CT-imaged nodules with subsequent surgical resection and storage as formalin-fixed paraffin-embedded tissue. CT images were reviewed by 2 observers and classified as pure non-solid or solid nodules on CT attenuation. The corresponding surgical resections were reviewed for pathology, and invasion was measured using Aperio whole scanned slides. Faxitron x-ray of the paraffin blocks was used to identify the areas of lowest and highest tissue density for punch samples. Tissue microarrays slides of 3 ROIs (1 mm diameter punch cores) per samples were generated. Normal tissue was obtained from separate paraffin blocks of tissue taken away from the tumor mass. The manual tissue arrayer (Beecher Instruments, Sun Prairie, WI) was used to generate tissue microarray (TMA) slides.

### Multiplex imaging

The immunofluorescence imaging was performed using the Neogenomics platform as previously described (11). Briefly, formalin-fixed paraffin-embedded tissue arrays were baked at 65°C for 1 h. Slides were deparaffinized with xylene, rehydrated by decreasing ethanol concentration washes, and then processed for antigen retrieval. A two-step antigen retrieval was adopted to allow antibodies with different antigen retrieval conditions to be used together on the same samples (12). Samples were then blocked against nonspecific binding with 10% (wt/vol) donkey serum and 3% (wt/vol) bovine serum albumin (BSA) in phosphate-buffered solution (PBS) for 1 h at room temperature and stained with DAPI for 15 min. Directly conjugated primary antibodies were diluted in PBS supplied with 3% (wt/vol) BSA to optimized concentrations and applied for 1 h at room temperature on a Leica Bond III Stainer. In the case of primary-secondary antibody staining, samples were incubated with primary antibody, followed by incubation with species-specific secondary antibodies conjugated to either Cyanine 3 (cy3) or cyanine 5 (cy5).

Stained images were collected on INCell analyzer 2200 microscope (GE Healthcare Life Sciences) equipped with high-efficiency fluorochrome specific filter sets for DAPI, cy3 and cy5 (11). For multiplexed staining where co-localization was desired, the regions of interest (∼0.4– 0.6 mm^2^ tissue area) were imaged, and stage coordinates were saved. The coordinates of each image region were then recalled for each subsequent round after minor readjustment using reference points from the first-round DAPI image and determining the appropriate offset. The exposure times were set at a fixed value for all images of a given marker. For image analyses, microscopy images were exported as full-resolution TIFF images in grayscale for each individual channel collected.

### MultiOmyx image analytics

The acquired images from sequential rounds were registered using DAPI images acquired in the first round of staining via a rigid registration algorithm for each region of interest. The parameters of transformation were then applied to the subsequent rounds, which ensured that the pixel coordinates across all the imaging rounds corresponded to the same physical locations on the tissue. Classification and co-expression analysis were performed in multiple stages. First, a nuclear segmentation algorithm was applied on the DAPI image to delineate and identify individual cells. Location information and expression of all the markers were computed for every cell identified. Then, morphologic image analysis and shape detection were performed using proprietary algorithms. These algorithms detect and classify cells as positive or negative for each marker depending on their subcellular localization and morphology. A tissue-quality algorithm was also applied to the images to ensure image artifacts that arose owing to tissue folding or tear did not affect cell classification. Co-expression analysis and phenotype identification were performed by combining individual marker classification results.

### αSMA Morphology

The αSMA morphology of 3 ROIs per tumor were independently scored by 2 investigators (NKA and TEM) as either organized (value of 1) or disorganized (value of 0). Representative images of these 2 classes of morphology are shown in Figure 9. Each tumor was assigned a predominant αSMA morphology based on the sum score: disorganized ≤ 1; organized ≥ 2.

### Statistical Analysis

Students unpaired T tests or Fishers exact test, where appropriate, were used to evaluate group differences (Prism, GraphPad Software). All reported p values are significant based on Benjamini-Hochberg control for false discovery at an alpha of 0.05.

## RESULTS

### Defining the tumor microenvironment by multiplex immunofluorescence analyses

We used multiplex immunofluorescence microscopy to contrast the TME of both pure non-solid CT scan density (p-NS) and solid early stage human LUADs. We generated a tissue microarray (TMA) of stage 1A LUADs: 25 p-NS tumors and 27 solid tumors (supplemental Table I). To capture the heterogeneity within individual tumors the TMA was constructed from different cores (regions of interests, ROI’s) of each tumor. The expressions of all markers were determined in at least 3 ROIs per tumor and some markers were measured in 6 ROIs per tumor. A TMA of normal lung was generated from tissue adjacent to the tumors of 49 subjects (a single core per sample). The Neogenomics multiplex immunofluorescence platform (11) was used to quantify expressions of 19 markers to define the immune cell composition and activation states, to identify cancer/epithelial cells and fibroblasts (Fig. 1A and supplementalTable II). Briefly, the TMAs are stained with two different antibodies directly labeled with either Cy3 or Cy5 fluorophores. Images in both channels are collected, the Cy3 and Cy5 fluorescence inactivated, samples washed, stained with 2 new antibodies directly labeled with Cy3 or Cy5, images collected and the process repeated (12). In this iterative fashion the TMAs were stained with 20 antibodies. A representative multiplexed image is shown in figure 1B.

**Figure 1.**
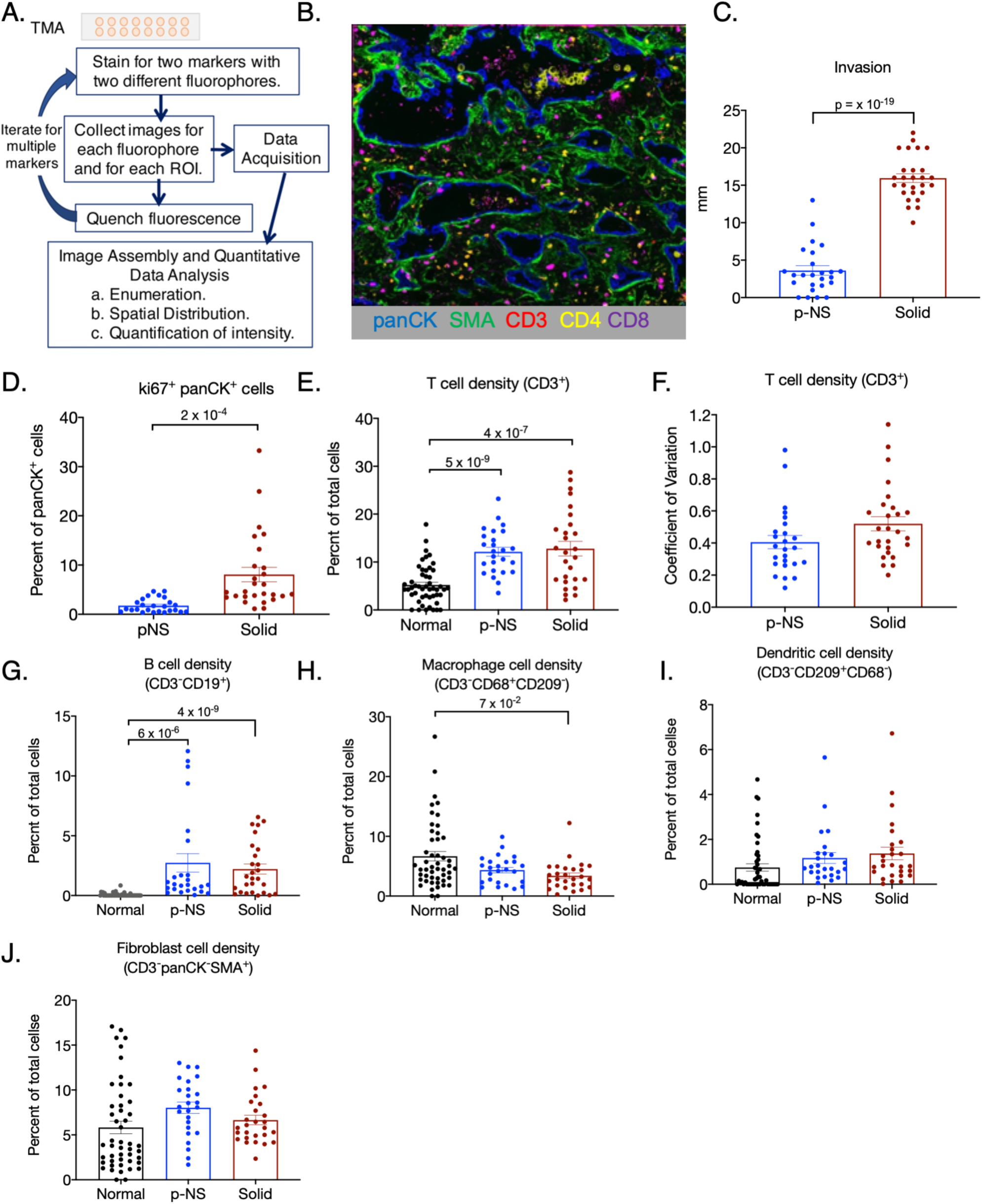
Immunofluorescence immune profiling. **A**. Cartoon of Neogenomics iterative, quantitative immunofluorescence platform. **B**. Representative Neogenomics pseudo-colored multiplex image, 200x magnification. Each channel scaled for purpose of presentation. **C**. Cancer cell invasion of the p-NS and solid tumor groups. **D**. Percent of panCK^+^ cells that are ki67^+^. **E**. Density of CD3^+^ cells determined as the percentage of total cells per ROI. **F** Coefficient of variations of CD3^+^ cell densities of the 6 ROIs per tumor of the p-NS and solid groups. **G**. Density of B cells (CD3^-^CD19^+^ cells) determined as the percentage of total cells per ROI. **H**. Density of macrophages (CD68^+^CD3^-^ CD209^-^ cells) determined as the percentage of total cells per ROI. **I**. Density of dendritic cells (CD209^+^CD3^-^CD68^-^ cells determined as the percentage of total cells per ROI. **J**. Density of fibroblasts (CD3^-^ panCK^-^ SMA^+^) determined as the percentage of total cells per ROI. In all panels the symbols for normal tissue are data from a single ROI per sample, whereas in the p-NS and solid groups each symbol is the mean of 3 to 6 ROIs per tumor. Total cells were determined by DAPI stained nuclei per ROI. The group means ± SEM are shown. Statistically significant differences are noted. Student’s t tests, unpaired. All p values shown are significant based on Benjamini-Hochberg control for false discovery at an alpha of 0.05.

The CT density-defined groups were matched for age, sex and tumor size (supplemental Table I). In the p-NS group there were equal numbers of EGF receptor (EGFR) mutant and KRAS tumors (28% each), whereas in the solid group there were 3 times as many KRAS (37%) as mutant EGFR tumors (11%). Within the constraints of our sample size, we were unable to detect a significant differences in the immune compositions of KRAS and mutant EGFR TMEs, either within a tumor group or between the two tumor groups. Tumors in the CT solid group were significantly more invasive than the p-NS tumors (Fig. 1C), supporting the use of CT density as a means to classify tumors for analyses of how the TME changes with tumor progression. The percentage of Ki67^+^ panCK cells in the solid tumor group was significantly increased relative to the p-NS, confirming that the CT dense tumors are more proliferative (Fig. 1D).

### p-NS and solid tumors are equally T cell inflamed compared to normal lung

T cell density (CD3^+^ cells) was significantly elevated in both the p-NS and solid tumor groups compared to the normal lung group, whereas there was no significant difference in T cell density between p-NS and solid tumor groups (Fig. 1E). Therefore, neither the emergence of a p-NS tumor from normal lung nor the progression from p-NS to solid tumors are characterized by exclusion of T cells from the tumor mass. In addition, intra-tumor heterogeneity in T cell densities, assessed by the coefficient of variation of T cell density among the 6 ROIs per tumor, were similar in the two tumor groups, establishing that intra-tumor heterogeneity in T cell density does not change significantly during progression of p-NS to solid tumors (Fig. 1F).

To broadly define the TME of the groups, we also determined the density of B cells (CD3^-^CD19^+^ cells), macrophages (CD3^-^ CD68^+^ CD209^-^ cells), dendritic cells (CD3^-^ CD209^+^ CD68^-^ cells) and fibroblasts (CD3^-^ panCK^-^ αSMA^+^ cells). Alpha smooth muscle actin (αSMA) is frequently used as a marker of cancer-associated fibroblasts, although it is expressed in other cell types, most prominently myofibroblasts and pericytes (13). Here we refer to αSMA^+^ cells as fibroblasts. B cell density in the normal lung was low (0.06% of total cells) and significantly elevated in both p-NS and solid tumor groups (2.7% and 2.1% of total cells, respectively) (Fig. 1G). However, there was no significant difference in B cell density between the two tumor groups. There was a small yet significant reduction in macrophage cell density in the solid tumor groups compared to normal lung, whereas there was no difference between the p-NS group and the normal lung (Fig. 1H). Neither the dendritic cell density nor fibroblast cell density varied significantly among the three groups (Fig. 1I & J).

### The TMEs of p-NS tumors are enriched for Treg cells

We investigated the T cell subtype compositions to determine whether the TME of p-NS tumors were predominantly immune-suppressive. CD4^+^ T cells (CD3^+^CD4^+^) were significantly increased in the p-NS tumor group (Fig. 2A). The subtype composition of the CD4^+^ T cells also varied between the normal lung and p-NS tumor groups. Immune-suppressive Treg cells (CD3^+^ CD4^+^ FoxP3^+^ cells) were significantly increased in the p-NS tumor group at the expense of T helper cells (CD3^+^ CD4^+^ FoxP3^-^ cells) and approximately 5% of the Treg cells in the p-NS group were Ki67^+^ (Fig. 2B&C). Importantly, CTLA-4^+^ Treg cells were enriched in the p-NS tumor group compared to normal lung, approximately 2% of which were Ki67^+^ (Fig. 2D&E). CTLA-4 expression is associated with enhanced Treg activity and homeostasis (14,15). These data demonstrate an increased immune-suppressive TME in p-NS tumor group compared to normal lung.

**Figure 2.**
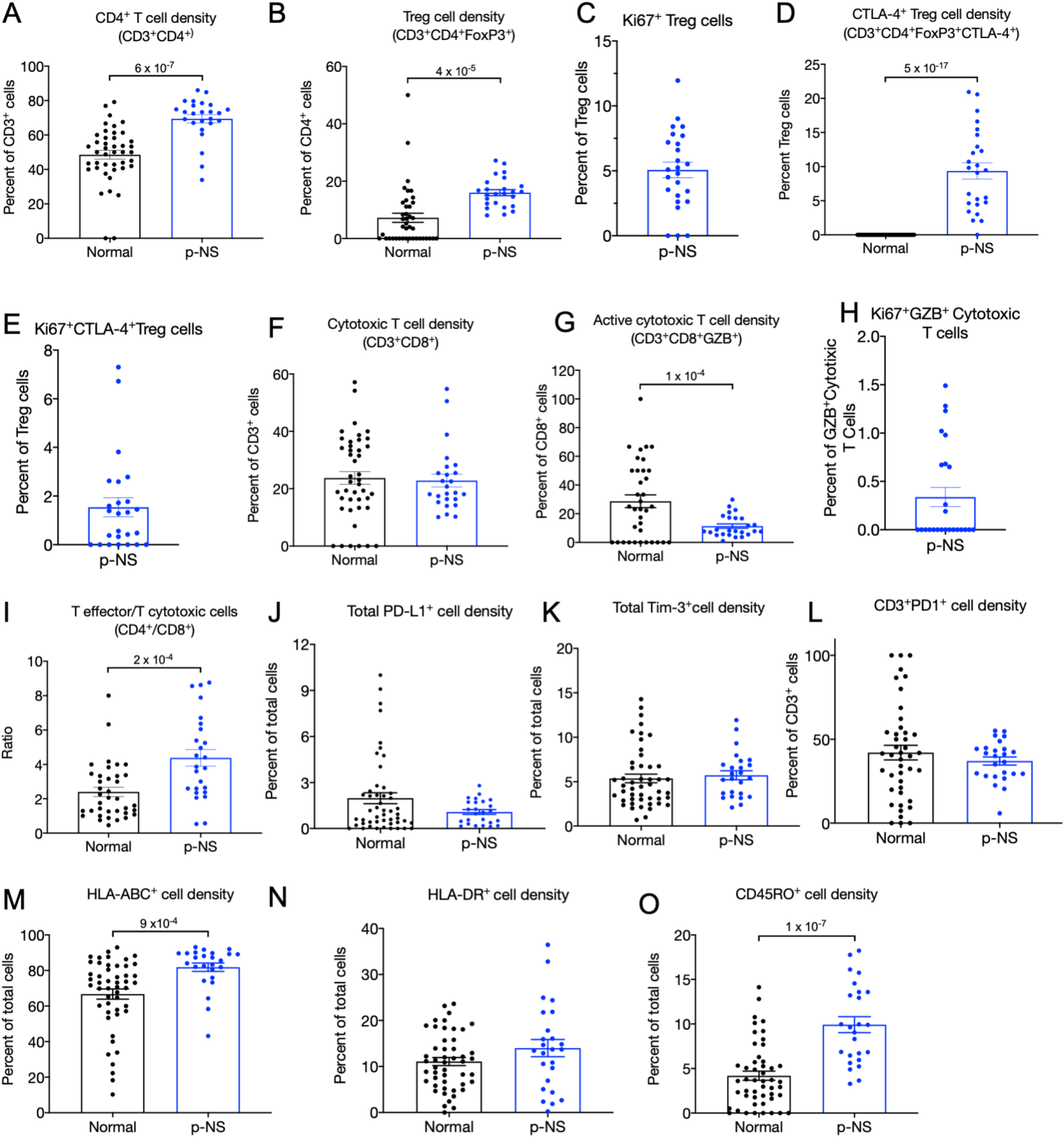
T cell composition. **A**. Densities of CD4^+^ cells determined as percentage of CD3^+^ cells. **B**. Treg cell densities (CD3^+^CD4^+^FoxP3^+^) determined as the percentage of CD3^+^CD4^+^ cells. **C**. Percent of Tregs in p-NS tumor group that are Ki67^+^. **D**. CTLA-4^+^ Treg cell densities (CD3^+^CD4^+^FoxP3^+^CTLA-4^+^) as the percentage of CD3^+^CD4^+^FoxP3^+^ cells. **E**. Percent of CTLA-4^+^ Tregs in p-NS tumor group that are Ki67^+^. **F**. Cytotoxic T cell densities (CD3+CD8^+^) determined as the percentage of CD3^+^ cells. **G**. Active cytotoxic T cells densities (CD3^+^CD8^+^GZB^+^) determined as the percentage of CD3^+^CD8^+^ cells. GZB, granzyme B. **H**. Percent of active cytotoxic T cells in p-NS tumor group that are Ki67^+^. **I**. Ratios per tumor of Treg cells (CD3^+^CD4^+^FoxP3^+^) to cytotoxic T cells (CD3+CD8^+^). **J**. Total PD-L1^+^ cell densities as a percentage of total cells. **K**. Total Tim-3^+^ cell densities determined as a percentage of total cells per ROI. **L**. Total PD1^+^ cells densities determined as a percentage of total cells per ROI. **M**. Total HLA-ABC^+^ cell densities determined as a percent of total cells per ROI. **N**. Total HLA-DR^+^ cell densities determined as a percentage of total cells per ROI. **O**. Total CD45RO^+^ cell densities determined as a percentage of total cells per ROI. In each comparison total cell numbers are defined by the number of DAPI stained nuclei per ROI. In all panels each normal tissue symbol is the cell density from a single ROI per sample, whereas in the p-NS group each symbol is the mean of 3 to 6 ROIs per tumor. The means ± SEM are shown for each group. Statistically significant differences are noted. Student’s T test, unpaired. All p values shown are significant based on Benjamini-Hochberg control for false discovery at an alpha of 0.05.

The density of cytotoxic T (CD3^+^CD8^+^) cells, as the proportion of T cells, did not differ between normal lung and p-NS tumor groups (Fig. 2F). However, granzyme B (GZB) expressing cytotoxic cells (GZB^+^CD8^+^CD3^+^ cells) were reduced in the p-NS tumor group relative to normal lung (Fig. 2G). GZB expression distinguishes active from resting or memory cytotoxic T cells. In addition, less than 1% of the active cytotoxic T cells in the p-NS tumor group were positive for the proliferation marker Ki67 (Fig. 2H). The ratio of Treg cells to cytotoxic T cells per individual sample was significantly elevated in the p-NS group that the normal lung group (Fig. 2I), supporting the hypothesis that in individual p-NS tumors both an increase in Tregs and a decrease in CD8 T cells contribute to an immune-suppressive TME.

Expression of immune checkpoints is a mechanism by which tumors evade immune-mediated killing (16). There was no difference between normal lung and p-NS tumor groups in two immune checkpoints known to be upregulated in LUADs (13): PD-L1^+^ or TIM-3^+^ (Fig. 2J&K). In addition, there was no difference in PD1^+^ T cells, whose expression is often used as marker of T cell exhaustion/dysfunction (17) (Fig. 2L). MHC class I^+^ cell density (HLA-ABC^+^) was elevated in the p-NS tumors, demonstrating that reduced MHC class I antigen presentation is not a mechanism for immune escape in p-NS tumors (Fig. 2M). There was no difference between normal lung and p-NS tumor group in MHC class II expression (HLA-DR^+^ cells) (Fig. 2N). However, there was an increase in the p-NS tumor group of CD45RO^+^ cells, a marker of memory T cells, consistent with an increase in antigen-experienced cells in the p-NS tumors (Fig. 2O).

Our analyses of the immune TME of p-NS tumors compared to adjacent normal lung support the hypothesis that in the p-NS tumors, an early manifestation of LUAD, cancer cells evade immune-mediated cell killing predominately through mechanisms involving a TME enriched with immune-suppressive Treg cells and depleted for activated cytotoxic T cells.

### Increased active Treg cells and cytotoxic T cells in the solid tumor group

We next compared the immune-TME of the p-NS and solid tumor groups by assessing the composition and activation states of the T cells. There was no difference in CD3^+^CD4^+^ cell density between p-NS and solid tumor groups, regardless of whether CD4^+^ cells were analyzed as the percent of total cells or as the percent of CD3^+^ cells (Fig. 3A & B). In both tumor groups about 70% of the T cells were CD4^+^ cells. However, the proportions of CD4^+^ cells that were immune-suppressive Treg cells were significantly increased in the solid tumor group (Fig. 3C). An equal proportion of Tregs were Ki67^+^ in both groups (Fig. 3D). However, the proportion of CTLA-4^+^ Treg cells was significantly elevated in the solid tumor group, consistent with an increased immune-suppressive TME of solid tumors relative to the p-NS tumors (Fig. 3E).

There was no significant difference in cytotoxic T cell (CD3^+^CD8^+^) density between the groups when measured as a proportion of total cells (density in tissue) or as a proportion of CD3^+^ cells (Fig. 3F&G). There was, unexpectedly, an increase in activated cytotoxic T cells (CD3^+^CD8^+^GZB^+^) as well as Ki67^+^ active CD8 cells in the solid tumor group (Fig. 3H&I). Thus, there is an active cytotoxic immune response in the more advanced solid tumors. The corresponding increase in immune-suppressive Treg cells may serve to counter-balance the increase in cytotoxic cells, thereby contributing to a blunting of activated cytotoxic T cells. Supporting that hypothesis, there was a positive correlation between the densities of Treg cells and GZB^+^CD8 cells in the solid tumor group (Fig. 3J).

**Figure 3.**
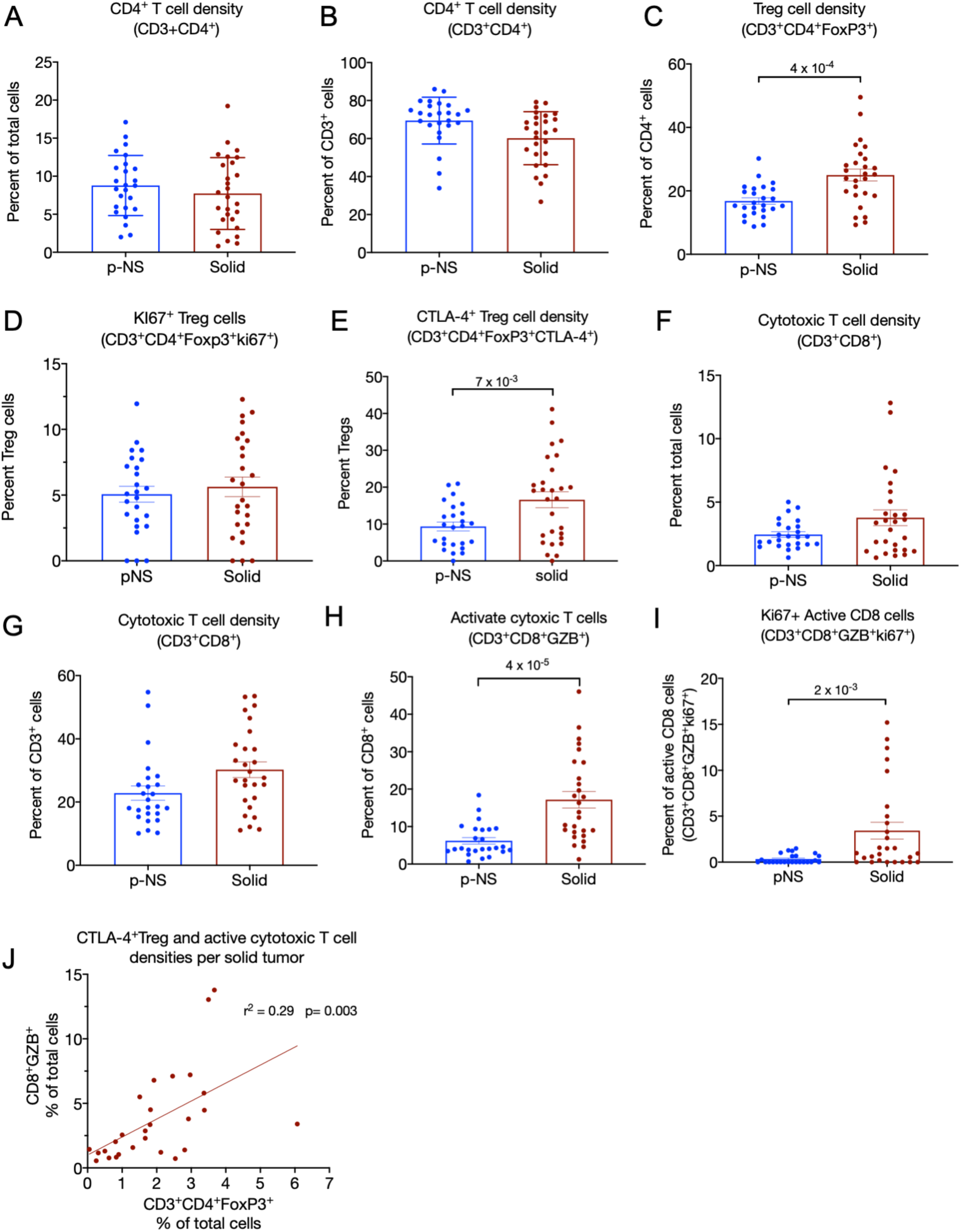
T cell compositions of p-NS and Solid tumors. **A**. CD4^+^ cell densities determined as a percentage of total cells per ROI. **B**. CD4^+^ cell densities determined as a percentage of CD3^+^ cells. **C**. Treg cell densities (CD3^+^CD4^+^FoxP3^+^) determined as a percentage of CD4^+^ cells. D. ki67^+^ Treg cell densities (CD3^+^CD4^+^FoxP3^+^ki67^+^) determined as a percentage of Treg cells. **E**. CTLA-4^+^ Treg cell densities (CD3^+^CD4^+^FoxP3^+^CTLA-4^+^) determined as a percentage of Treg cells. **F**. Cytotoxic T cell densities (CD3^+^CD8^+^) determined as a percentage of total cells per ROI. **G**. Cytotoxic T cell densities (CD3^+^CD8^+^) determined as a percentage of CD3^+^ cells. **H**. Active cytotoxic T cell densities (CD3^+^CD8^+^GZB^+^) determined as a percentage of CD3^+^CD8^+^ cells. GZB, granzyme B. **I**. ki67^+^ active CD8+ cell (CD3^+^CD8^+^GZB^+^ki67^+^) densities determined as a percentage of CD3^+^CD8^+^ cells. **J**. Correlation of CTLA-4^+^ Treg (CD3^+^CD4^+^FoxP3^+^CTLA-4^+^) and active cytotoxic T cell (CD3^+^CD8^+^GZB^+^) densities as percentages of total cells per ROI. The p value is for the difference in the slope of the correlation line from 0. In each panel the symbols are the means of 3 to 6 ROIs per tumor. The means ± SEM are shown for each group. Statistically significant differences are noted. Students T test, unpaired. All p values shown are significant based on Benjamini-Hochberg control for false discovery at an alpha of 0.05.

We explored other potential immune-suppressive mechanism to account for the suppression of cytotoxic T cell response in the solid tumor group. There were no differences between the p-NS and solid tumor groups in PD1^+^ checkpoint expression in cytotoxic T cells (PD1^+^CD8^+^) (Fig. 4A). Hence, cytotoxic T cell “exhaustion” does not underlie immune suppression of the TME in solid tumors. There was, however, an increase in PD1^+^ Thelper cells (PD1^+^CD3^+^CD4^+^FoxP3^-^), indicating potential checkpoint-mediated blunting of T helper function contributing to the immune suppressive TME of solid tumors (Fig. 4B). There was no difference in expression of the TIM3^+^ checkpoint between the groups (Fig. 4C).

**Figure 4.**
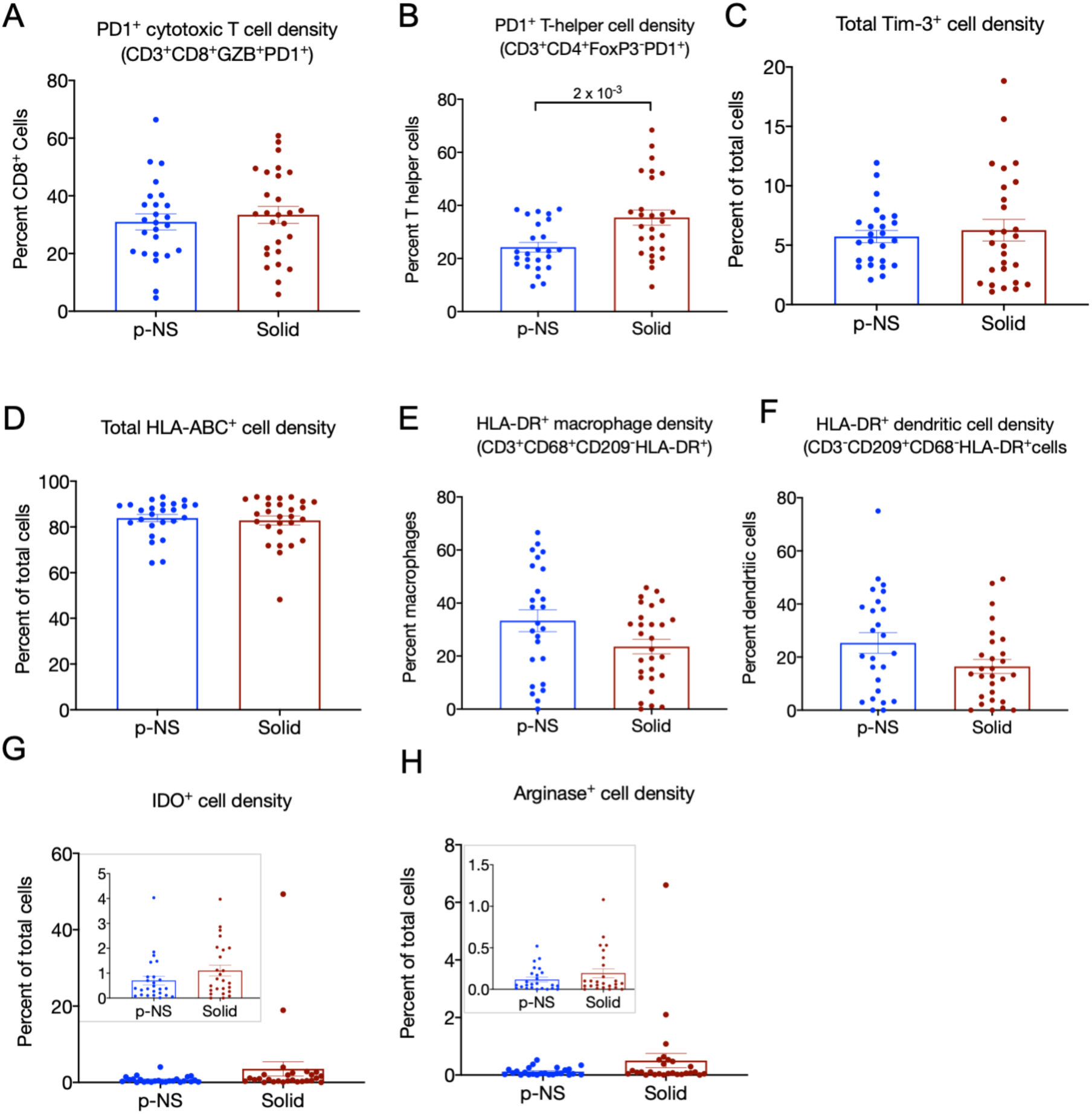
T cell activation states. **A**. PD1^+^ cytotoxic T cell densities determined as a percentage of CD8+ cells. **B**. PD1^+^ T helper cell (CD3^+^CD4^+^FoxP3^-^) densities determined as a percentage of T helper cells. **C**. Total TIM3^+^ cell densities. **D**. Total HLA-ABC^+^ cell densities determined as a percentage of total cells per ROI. **E**. HLA-DR^+^ macrophage densities determined as a percentage of total macrophages (CD3^-^CD68^+^CD209^-^). **F**. HLA-DR^+^ dendritic cell densities as percentage of dendritic cells (CD3^-^CD209^+^CD68^-^). **G**. IDO^+^ cell densities determined as a percentage of total cells. Inset, y axis expanded. **H**. Arginase^+^ cell densities determined as a percentage of total cells. Inset, y axis expanded. In all panels each symbol is the mean of 3 ROIs per tumor. The means ± SEM are shown for each group. Total cells determined by DAPI^+^ nuclei. Statistically significant differences are noted. Students T test, unpaired. All p values shown are significant based on Benjamini-Hochberg control for false discovery at an alpha of 0.05.

The densities of MHC class I-expressing cells (HLA-ABC^+^) were not different between the two tumor groups, demonstrating that in the more advanced solid tumors, cancer cells do not evade immune surveillance by reduced MHC class I antigen presentation (Fig. 4D). There were no differences in MHC class II expressing macrophages (HLA-DR^+^CD68^+^CD3^-^CD209^-^) nor dendritic cells (HLA-DR^+^ CD209^+^CD3^-^CD68^-^) between the groups (Fig. 4E&F).

Upregulation of the tryptophan catabolic enzyme, indoleamine 2,3-dioxyhydrogenase (IDO), can have an immune suppressive role in cancer (16). Because IDO can be expressed by either cancer cells or cells of the tumor stroma, we compared the p-NS and solid tumor groups for total IDO expressing cells (Fig. 4G). There was no difference in IDO expression between the tumor groups, although two solid tumors expressed IDO at a higher frequency than the others. Although IDO expression might have an immune suppressive role in these two solid tumors, increased IDO expression does not underlie enhanced immune suppression in the solid tumor group compared to the p-NS tumor group (inset Fig. 4G).

Arginase 1, expressed by cells of the TME, has an inhibitory effect on antigen-specific anti-tumor T cell responses (18). As was the case for IDO, Arginase expression does not distinguish the tumor groups, therefore Arginase I upregulation is not a common immune-suppressive mechanism in the solid tumor group (Fig. 4H). The frequency of arginase expression is elevated in 2 of the solid tumors (not the same as those in which IDO is upregulated), suggesting elevated Arginase might in some tumors have an immune suppressive role.

### Heterogenous expression of PD-L1 among the solid tumor group

Expression of the PD-L1 checkpoint is a mechanism for immune escape in NSCLC and the percent of cancer cells expressing PD-L1 is used clinically to assess overall PD-L1 expression in tumors (19). In the p-NS tumor group only 3 of 25 tumors had greater than 1% PD-L1^+^ panCK cells, whereas in the solid tumor group 13 of 29 had greater than 1% PD-L1+ panCK cells (Fig. 5A&B). Because of the broad range of PD-L1 expression among the solid tumors, we segregated them into three groups: tumors with less than 1% PD-L1^+^ panCK cells; tumors with less than 50% PD-L1^+^ panCK cells; tumors with greater than 50% PD-L1^+^ panCK cells. Total tumor-mass infiltrating T cells did not correlate with PD-L1 expression but higher PD-L1 expression in panCK cells correlated with an increase in activated cytotoxic T cells (CD3^+^CD8^+^GZB^+^) and CTLA-4^+^Tregs (CD3^+^CD4^+^FoxP3^+^) (Fig. 5C-E). No differences in the immune profiles of the low and intermediate PD-L1 expressing tumor groups were detected. Thus, a high percentage of PD-L1 expressing panCK^+^ cells was linked to increases in both immune suppressive and cytotoxic T cells. These data support the hypothesis that an anti-cancer cytotoxic T cell immune response was blunted by both the immune suppressive activities of Treg cells and an increase PD-L1 expression.

**Figure 5.**
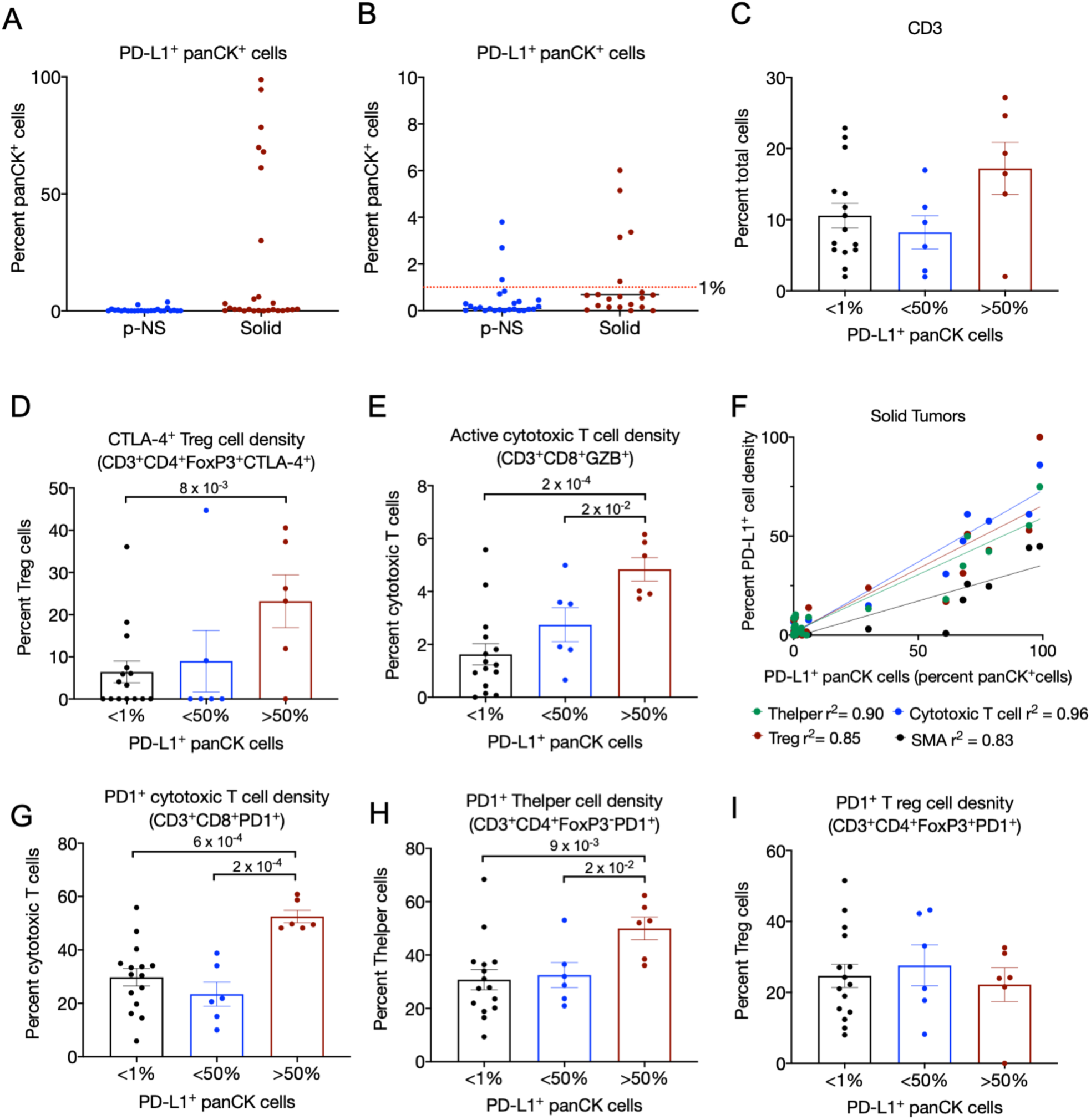
Immune profile linked to PD-L1 expression. **A**. PD-L1^+^panCK^+^ cells as a percentage of panCK^+^ cells. **B**. PD-L1^+^panCK^+^ cells, percent of panCK^+^ cells, on expanded y-axis to visualize differences among the low PD-L1 expressing tumors. The red dotted line is at 1% PD-L1^+^panCK cells. **C**., **D**., **& E**. Densities of CD3^+^ cells, CTLA-4^+^ Treg cells (CD3^+^CD4^+^FoxP3^+^CTLA-4^+^) and active cytotoxic T cells (CD3^+^CD8^+^GZB^+^) in the 3 subcategories of solid tumors segregated by panCK^+^ PD-L1^+^ expression, respectively. **F**. Correlation of panCK PD-L1^+^ expression and PD-L1 expression in Treg cells (CD3^+^CD4^+^FoxP3^+^), T helper cells (CD3^+^CD4^+^FoxP3^-^), cytotoxic T cells (CD3^+^CD8^+^) and αSMA^+^ cells. Correlation values (r^2^) are shown. **G**., **H**., and **I**. PD1^+^ Thelper cells (CD3^+^CD4^+^FoxP3^-^PD1^+^), PD1^+^ cytotoxic T cells (CD3^+^CD8^+^PD1^+^) and PD1^+^ Treg cells (CD3^+^CD4^+^FoxP3^+^PD1^+^) in the 3 subcategories of solid tumors segregated by panCK^+^ PD-L1^+^ expression, respectively. In all panels each symbol is the mean of 3 ROIs per tumor. The means ± SEM are shown for each group. Students T test, unpaired. Statistically significant differences are noted. All p values shown are significant based on Benjamini-Hochberg control for false discovery at an alpha of 0.05.

There was a positive correlation between PD-L1 expression in panCK^+^ cells and PD-L1 expression in T cells as well as fibroblasts within the same tumors (Fig. 5F). Thus, upregulation of PD-L1 is not restricted to cancer cells (panCK^+^ cells) but rather enhanced PD-L1 expression reflects conditions within the TME.

Expression of PD1 was increased in cytotoxic T cells of the high PD-L1 expressing tumor group (Fig. 5G). PD1 expression was increased in T helper cells of tumors in the highest PD-L1 expressing group compared to the intermediate and no PD-L1 expressing tumor groups (Fig. 5H). There were no differences among the groups in PD1^+^ Tregs (Fig. 5I). PD1, a negative feedback receptor, is expressed on both activated and “exhausted” T cells, and therefore its expression alone does not establish the activated state of T cells. However, high PD1 expression in an environment of high PD-L1 expression indicates active checkpoint function. Consequently, in the high PD-L1 expressing tumor group, both the anti-tumor T helper and cytotoxic T cells are likely to be suppressed, relative to the lower PD-L1 expressing tumor groups.

### T Cell exclusion from cancer cell nests as an immune-suppressive mechanism in p-NS tumors

The spatial distribution of T cells within the tumor mass, either localized to the stroma or infiltrated among the cancer cells nests, impacts immune response to the cancer cells since the physical proximity of T cells to cancer cells is required for cytotoxic T cell function. Restriction of intra-tumoral T cells to the stroma (fibroblast-rich) and exclusion from the parenchyma is a mechanism for evasion of immune surveillance (2,5,20,21).

We quantified the spatial distribution of T cells within the stroma or cancer cell nests based on a panCK^+^ mask. A frequency distribution histogram based on T cell infiltration of cancer cell beds (percent total tumor T cells) revealed increased T cell infiltration in the solid tumor group (Fig. 6A). To explore differences in the immune composition based on localization within the tumor (that is, stroma or cancer cell nests), we used a value of greater than or equal to 5% of T cells (CD3^+^) within cancer cells nest to segregate tumors into ‘high’ and ‘low’ T cell infiltration of cancer cell nests (Fig. 6A). There was a trend towards increased infiltration of T cells among the cancer cell nests in the solid tumor group: 74% of the solid tumors and 52% of the p-NS had high T cell infiltration of the cancer cell nests (p=0.15, Fischer’s Exact Test). However, differences in the compositions of the T cells within the cancer cells nests between the two tumor groups were similar to those differences in the tumor not segregated for location, with an increase of both Treg cells and cytotoxic cells in the solid tumor group (Fig. 6B&C).

**Figure 6.**
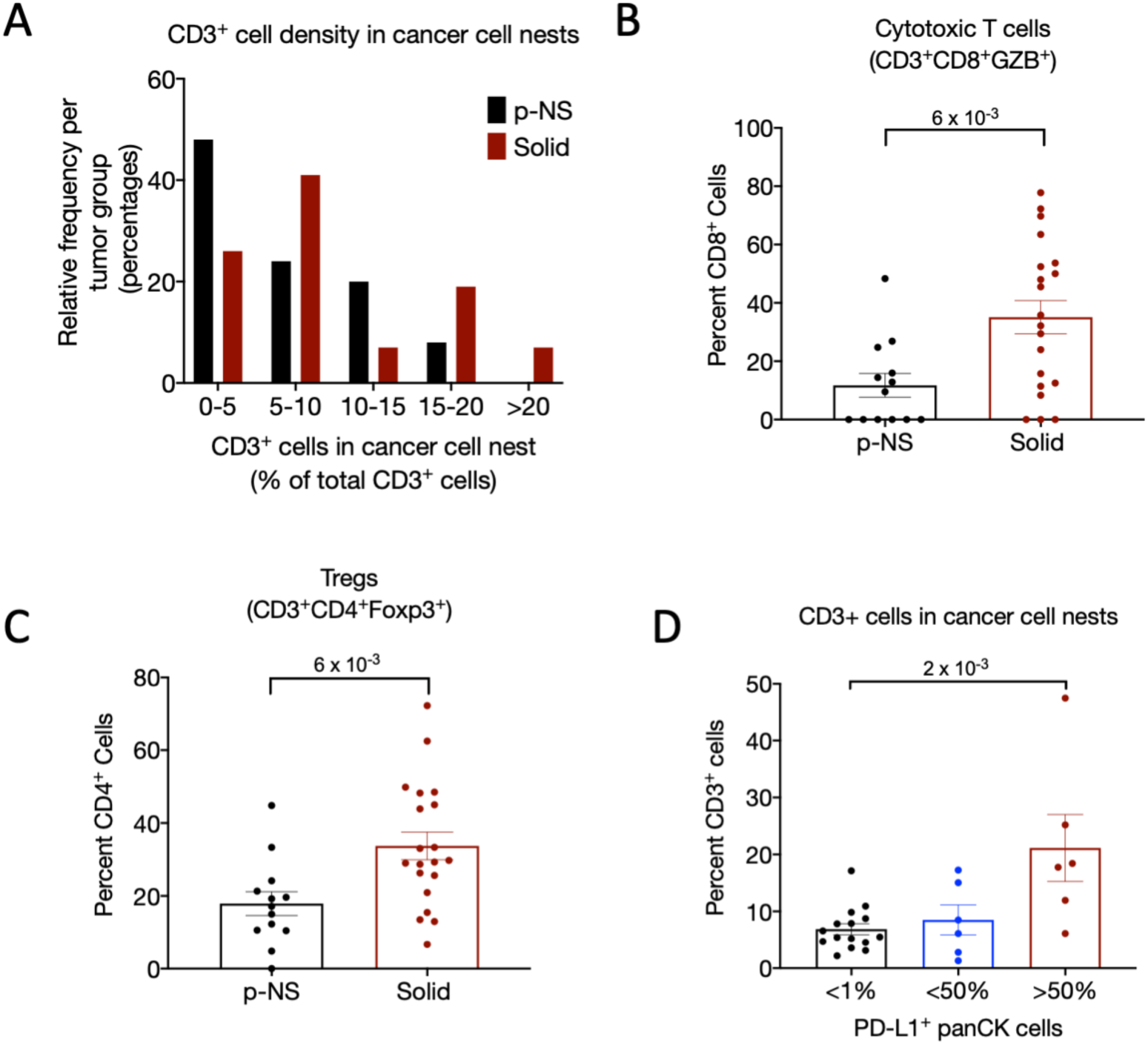
Distribution of T cells among cancer cell nests. **A**. Frequency distribution of p-NS and solid tumors based on percent of T cells located within the cancer cell nests. **B**. Cytotoxic T cells (CD3^+^CD8^+^) within the cancer cell nests as a percent of CD3^+^ cells in the tumor. **C**. Tregs (CD3^+^CD4^+^FoxP3^+^) within the cancer cell nests as a percent of CD4+ cells in the tumor. **D**. T cells (CD3^+^) within the cancer cell nests as a percentage of total T cells in the 3 subcategories of solid tumors segregated by panCK^+^ PD-L1^+^ expression. In all panels B,C & D, each symbol is the mean of 3 ROIs per tumor, and the means ± SEM are shown for each group. Students T test, unpaired. Statistically significant differences are noted. All p values shown are significant based on Benjamini-Hochberg control for false discovery at an alpha of 0.05.

T cell infiltration into the cancer cell nest of the solid tumors was highest in those tumors with the highest PD-L1 expression, in line with induced PD-L1 expression protecting the cancer cells from the cytotoxic T cells that had invaded the cancer cell nest (Fig. 6D). There were no differences in total tumor-mass infiltrating T cells between the p-NS and solid tumor groups (Fig. 1E). Therefore, exclusion of the T cells from the cancer cell nests, rather than from the tumor mass per se, is potentially a more important mechanism for immune escape in the p-NS tumor group than it may be in the solid tumor group.

### Fibroblasts morphology varies between the p-NS and solid tumors groups

Staining for alpha smooth muscle actin (*α*SMA) as a marker of fibroblasts revealed two prominent morphologies represented in both tumor groups. In one, the *α*SMA was well organized, circumscribing the cancer cell nests (panCK^+^) (Fig. 7A&B). In the other the *α*SMA pattern was disorganized and fragmented (Fig. 7C&D). The disorganized fibroblasts structure (*α*SMA morphology) was significantly more prevalent in the solid tumors (Fig. 7E). These data associate a disorganized *α*SMA morphology with the more advanced tumor group. Furthermore, in the p-NS tumor group the degree of T cell infiltration among the cancer cells was significantly decreased in tumor areas (ROIs) of organized *α*SMA morphology, correlating the intact fibroblastic structure with reduced accumulation of T cell among the cancer cells, a correlation not observed in the solid tumors (Fig. 7F).

**Figure 7.**
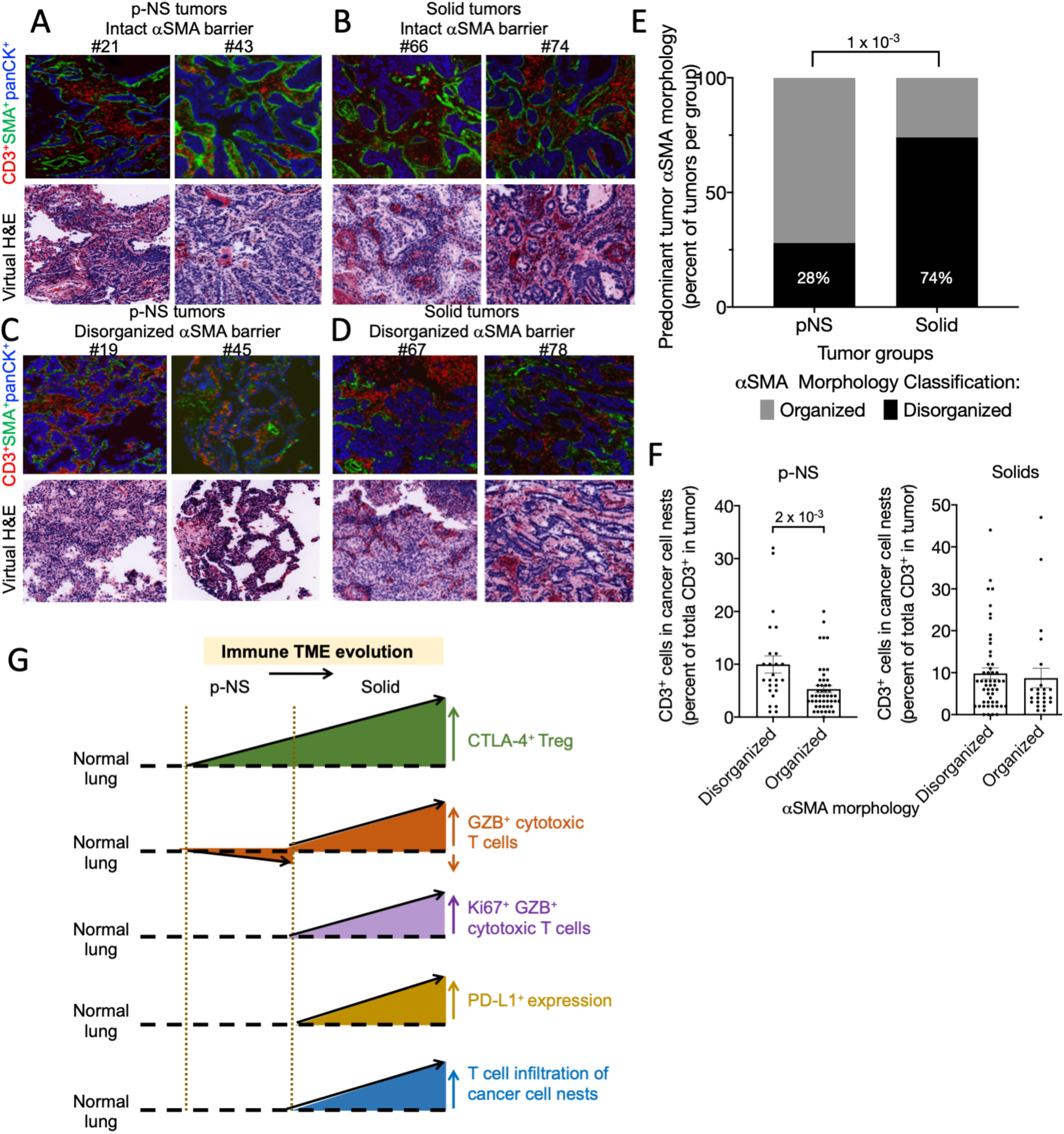
Relationships between tumor stroma and cancer cell nests. **A**. Immunofluorescence and corresponding virtual H&E of two p-NS tumors with an intact αSMA barrier, **B**. two solid tumors with an intact □SMA barrier; **C**. two p-NS tumors with a disrupted □SMA barrier; **D**. two p-NS tumors with a disrupted αSMA barrier. All images acquired at 200x magnification. **E**. Predominant tumor αSMA morphology by tumor group. p value, Fischer’s Exact test. **F**. CD3^+^ cell infiltration of cancer cell nest per ROI grouped by αSMA morphology of the ROI. Left are p-NS tumors and right Solid tumors. Statistically significant differences are noted. Unpaired Student’s T test. **G**. Model for evolution of immune TME is stage 1A NSCLC. See text for discussion. Vertical dotted lines are separation between p-NS and Solid groups.

## DISCUSSION

Immune escape is essential for tumor progression, and a more thorough understanding of the cellular and molecular mechanism(s) by which preinvasive malignancies escape immune surveillance may allow for the development of more effective interception strategies that can be deployed at the earliest manifestations of disease (20-22). Currently there are significant gaps in our understanding of the molecular and cellular events driving immune escape in preinvasive lung lesions. Although there have been recent reports on the genomic and transcriptomic landscapes of preinvasive squamous cell cancer (23,24) there are only a few studies of pre-invasive pulmonary adenocarcinoma, and even fewer that focused on the immune milieu in tumor microenvironment. The most comprehensive profiling of the TME, reported by Lavin *et al*, was focused on clinically evident early lung cancers rather than precursor lesions (25). A more recent report by Chen et al, reported on 98 precursor lesions of lung adenocarcinoma and found that immune infiltration is correlated with copy number alterations of chromosome arm 6p suggesting a link between arm-level events and the tumor immune environment (26). Our limited understanding of the early events associated with progression of precursor lesions of LUAD is at least partly due to the difficulty in accessing sufficient material from these small lesions in humans for study. Typically, LUAD precursor lesions are situated peripherally in the lung parenchyma where they are inaccessible by bronchoscopic approaches and therefore tissue sampling is only possible after surgical removal. In this study we contrasted two extremes on the radiographic spectrum, the pure non-solid nodule which generally (but not always) harbors pre or minimally invasive adenocarcinoma and the solid nodule representing frankly invasive cancers. The choice of extremes was designed to allow us to maximize and capture significant molecular events associated with transition to a more invasive phenotype.

### Evolution of preinvasive adenocarcinoma in pure non-solid tumors

We found that compared to adjacent normal lung, the stroma of p-NS tumors was intensely infiltrated by immune cells, an infiltration predominantly composed of CD3^+^ T cells. The composition of the CD3^+^ population was dominated by CD4^+^ cells that accounted for nearly two thirds of all T cells within the tumor nodule. Significantly, immune-suppressive regulatory T cells (CD3^+^CD4^+^FoxP3^+^) were twice as prevalent in the p-NS tumor group than in normal lung (Fig. 2). Approximately 10% of the Tregs in p-NS group expressed the activation marker CTLA-4, whereas less than 1% of Tregs in normal lung were CTLA-4^+^ (Fig. 2). Although, the proportion of cytotoxic T cells (CD3^+^CD8^+^) in the p-NS tumor nodule group was essentially the same as in normal lung, the proportion of GZB^+^ CD8^+^ T cells was almost three-fold lower than that in normal lung and nearly all lacked Ki67 expression, suggesting a possible exhausted phenotype (Fig. 2). On a per tumor basis, the Treg cell-to-CD8^+^ cell ratio was significantly higher in the tumor nodule than in normal lung, indicating the dominance of Treg-mediated immunosuppressive effect. We also found that the expression of MHC class I molecules and CD45RO^+^ cells were significantly higher in tumor nodules than in normal lung, demonstrating that escape from immune surveillance was not mediated by aberration in antigen presentation (Fig. 2). Interestingly, expression of the inhibitory checkpoints PD1, PD-L1 and Tim-3, were not upregulated in the p-NS tumor group, suggesting that immune privilege of p-NS nodules does not result from activation of inhibitory immune checkpoints.

### Evolution of early solid invasive adenocarcinoma

We made several observations when comparing the immune TME profile of lung adenocarcinoma presenting as a solid tumor with that arising in p-NS nodules. Our findings strongly support the hypothesis that multiple modes of tumor escape develop during transition to an invasive phenotype. First, although adenocarcinomas arising in both types of nodules were equally T cell inflamed (Fig. 1), those in solid nodules had a significantly higher proportion of regulatory T cells, with a two-fold increase in the proportion of immune suppressive Treg cells (CD3^+^CD4^+^FoxP3^+^) that expressed CTLA-4 (Fig. 3). There was also an associated three-fold increase in the number of activated cytotoxic T cells (GZB^+^CD8^+^) cells in the solid tumors (Fig. 3), suggesting a nascent albeit ineffective anti-tumor immune response. It is likely that any emergent anti-tumor immune response may have been at least partially restrained by Treg cell-mediated immune suppression. In the context of the immune editing hypothesis for tumor progression, these data indicate that the transition from p-NS to solid nodules reflects a shift from equilibrium to escape phase of immune editing (9). Second, while there was essentially no PD-L1^+^ expression in panCK^+^ cells of the p-NS group, collectively solid nodules had a significantly higher frequency of PD-L1^+^panCK^+^ cells (Fig. 5). Thus, expression of the PD-L1 inhibitory immune checkpoint, an established mode of immune evasion potentially deployed by cancer cells possibly due to the release of interferon gamma from activated cytotoxic T cells, was only observed in more advance tumors.

### PD-L1 expression defines two immune phenotypes of solid adenocarcinoma

PD-L1 expression in panCK^+^ adenocarcinoma presenting as a solid nodule identified three distinct sub-groups based on the frequency of PD-L1 expression. Tumors in which 50% or more of the panCK^+^ cells expressed PD-L1, a threshold commonly used in the clinic for treating stage IV lung cancer patients with anti-PD-1 alone (that is, without chemotherapy), were grouped as high PD-L1 expressers. Tumors in which greater than 1% but less than 50% of the panCK^+^ were PD-L1^+^ were classified as a separate group as were those in which PD-L1 expression in panCK^+^ was less than 1%. Tumors with high PDL-1 expression in panCK^+^ cells were associated with higher PD-L1 expression in various T cell subsets and fibroblasts. High expressers had a distinctive immune phenotype characterized by more intense T cell infiltration, and broad activation of the adaptive immune response, including significantly higher frequencies of PD1^+^ cytotoxic T cells (CD3^+^CD8^+^GZB^+^) and PD1^+^ cytotoxic T helper cells (CD3^+^CD4^+^ FoxP3^-^) and CTLA-4^+^ Treg cells (CD3^+^CD4^+^ FoxP3^+^). This defines tumors that are most likely to respond to dual immune check point inhibition targeting CTLA-4 and/or the PD1/PDL-1 axis. There was no significant difference in either the composition or activation state of immune cells of the TME between low and intermediate expressers of PD-L1, which might reflect a relatively quiescent immune response. In this group of patients, response to immune checkpoint inhibition alone may be unlikely and this is reflected to some extent in clinical practice where such patients are treated by a combination of chemotherapy and immune checkpoint inhibition (27).

### T cell infiltration into the cancer cell nests is also immune-suppressive

The phenomenon of T cell exclusion has been previously reported as a mechanism of immune evasion in multiple cancers including pancreatic, ovarian and colorectal cancer (28-30). A similar finding was observed in ductal carcinoma *in-situ* breast cancer where cancer cells are separated from tumor infiltrating T cells by a continuous layer of myoepithelial cells and myofibroblasts (31). T cell exclusion from the entire tumor mass was not observed in any of our cases. However, T cell exclusion from the cancer cell nests was observed in nearly 50% of all tumor nodules classified radiographically as p-NS and in 25% of solid nodules that represented frankly invasive adenocarcinoma. In all remaining cases there was infiltration of CD3^+^ T cells into the cancer cell nests where they were interspersed with the cancer cells. The immune phenotype of the T cells in the cancer cell nests mirrored that of CD3^+^ T cells in the stroma and was dominantly immune suppressive.

### Fibroblastic architecture possibly associated with T cell exclusion

To investigate whether fibroblastic spatial organization is associated with T cell exclusion we scored each tumor for continuity of αSMA staining. The fibroblastic barrier appeared continuous in 26% of all solid tumor nodules and interrupted or discontinuous in the remaining 74%. In the latter cases the CD3 cells were predominantly interspersed among the cancer cells. In contrast, in p-NS tumors the barrier was continuous in three quarters of patients and discontinuous in the remainder. We found that a continuous fibroblastic barrier was significantly associated with T cell exclusion in p-NS tumor group but not in the solid nodule group, probably because the majority of solid nodules had a discontinuous barrier. Whether T cell exclusion in our samples was the result of a mere physical structural separation or alternatively mediated by specific chemokine/receptor signaling or both remains to be elucidated. Fearon and co-workers have shown that ligation of CXCL12 expressed on cancer associated fibroblasts to its receptor CXCR4 on cancer cells may contribute to T cell exclusion in pancreatic adenocarcinoma (32).

In summary, our data suggest a complex dynamic interaction between the tumor and its immune microenvironment (Fig. 7G). A dominant regulatory T cell-mediated immune suppression is initiated at the precursor level and is sustained with rising intensity throughout malignant progression. T cell exclusion from the cancer cell nests appears to be an additional mechanism of immune evasion deployed by some early invasive lung cancers, possibly until tumor evolution leads to a durable, viable invasive phenotype that breaks down the fibroblastic barrier. Throughout the entire process a nascent effector immune response is present but is effectively thwarted by the immune-suppressive elements. These data suggest that different interception strategies should be employed at different stages of tumor evolution.

## Abbreviations

TME: tumor microenvironment
LUAD: lung adenocarcinoma
p-NS: Pure non-solid
AIS: adenocarcinoma *in-situ*
MIA: minimally invasive adenocarcinoma
GZB: granzyme B
ROI: region of interest
CT: computer tomography
EGFR: EGF receptor
TMA: tumor microenvironment

## Acknowledgements

The multiplex immunofluorescence was performed through a fee-for-service contract with Neogenomics Laboratories, Inc (Fort Myers, FL). The study was supported by UG3 CA244697 (NKA, ACB, VM, OE and TEM), the Yoram Cohen family foundation (NKA), Vicky and Jay Furhman family fund (NKA) and the WCM Meyer Cancer Center (NKA, TEM). In addition, OE is also supported by UL1TR002384, R01CA194547, LLS SCOR grants 180078-02, 7021-20. We thank Gary Koretzky (WCM), Niroshana Anandasabapathy (WCM) and Juan Cubillos-Ruiz (WCM) for helpful discussions and critical reading of the manuscript.

## Author contributions

**NKA:** conceptualized the project, contributed to experimental design, performed analysis and interpretations, and wrote the manuscript.

**ACB:** contributed to experimental design, data analyses, and edited the manuscript.

**VM:** contributed to experimental design and edited the manuscript.

**OE:** contributed to experimental design, data analyses, and edited the manuscript.

**TEM:** conceptualized the project, contributed to experimental design, performed analysis and interpretation, and wrote the manuscript.

---

**Supplemental Table I:**
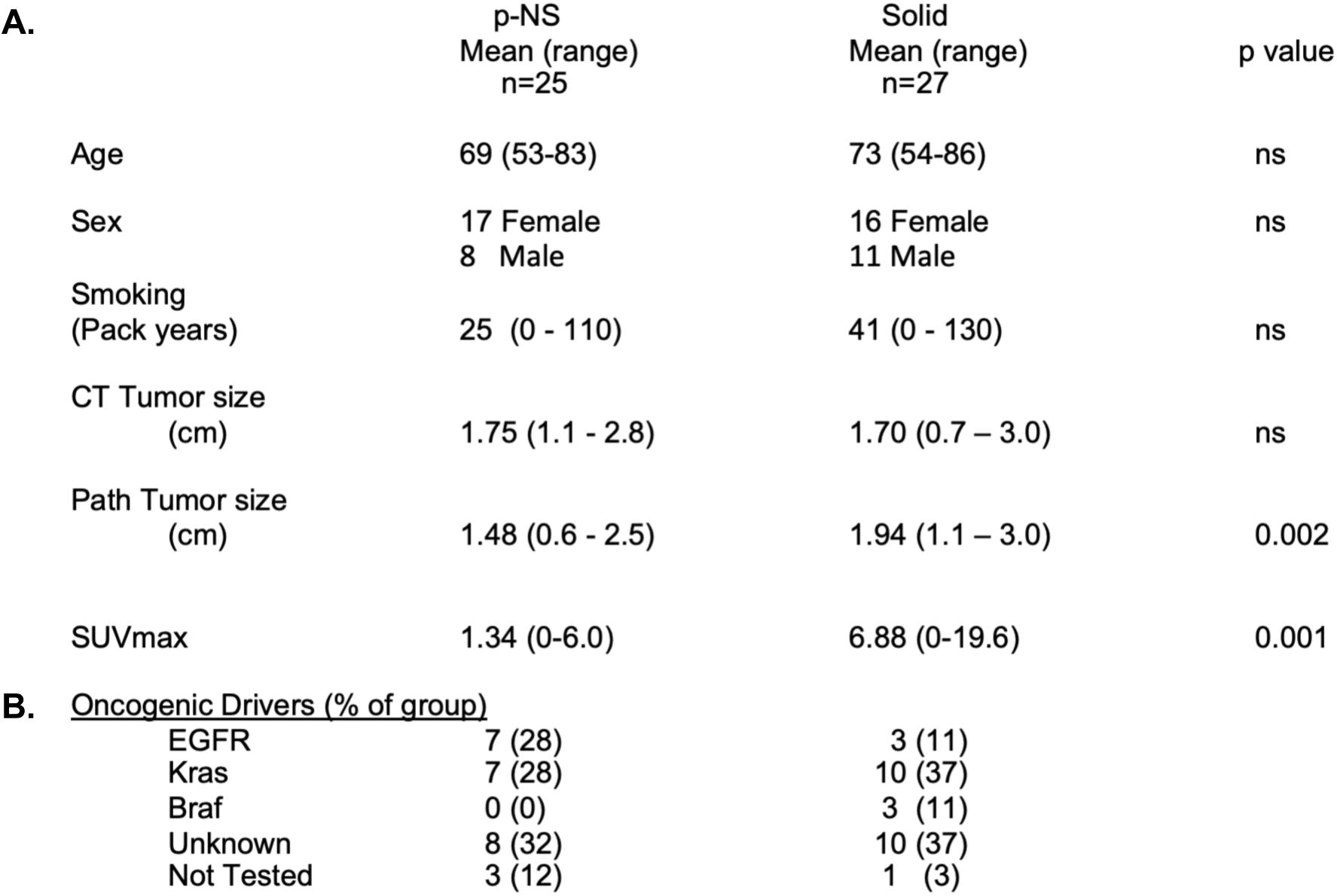
Clinical Characteristics.

**Table I. A. Subject profiles. B. Mutation status.**

**Supplemental Table II.**
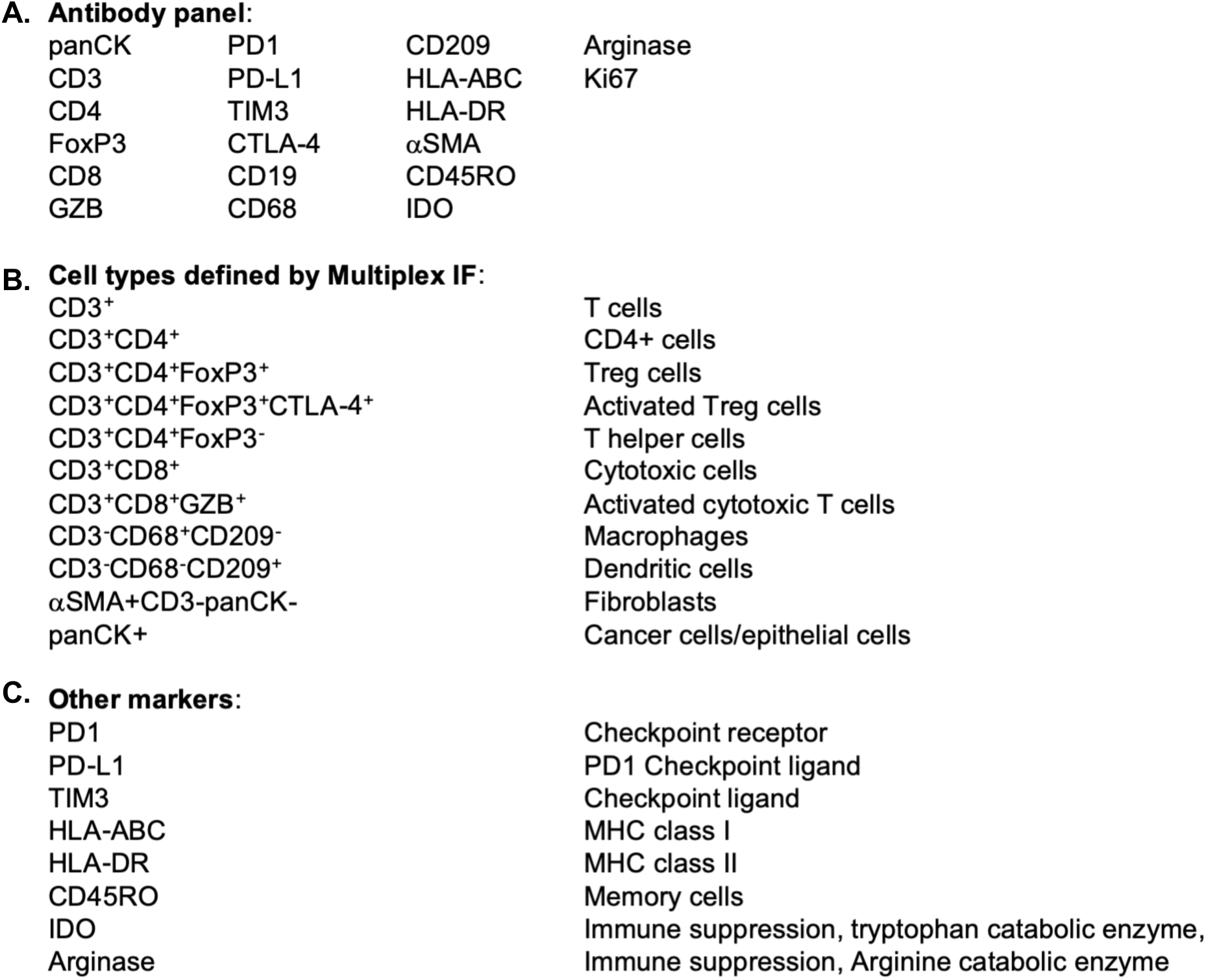
Profiling Markers.

A. Antibody panel:
|  |  |  |  |
| --- | --- | --- | --- |
| panCK | PD1 | CD209 | Arginase |
| CD3 | PD-L1 | HLA-ABC | Ki67 |
| CD4 | TIM3 | HLA-DR |  |
| FoxP3 | CTLA-4 | αSMA | |
| CD8 | CD19 | CD45RO |  |
| GZB | CD68 | IDO |  |

B. Cell types defined by Multiplex IF:
|  |  |
| --- | --- |
| CD3<sup>+</sup> | T cells |
| CD3<sup>+</sup>CD4<sup>+</sup> | CD4<sup>+</sup> cells |
| CD3<sup>+</sup>CD4<sup>+</sup>FoxP3<sup>+</sup> | Treg cells |
| CD3<sup>+</sup>CD4<sup>+</sup>FoxP3<sup>+</sup>CTLA-4<sup>+</sup> | Activated Treg cells |
| CD3<sup>+</sup>CD4<sup>+</sup>FoxP3<sup>-</sup> | T helper cells |
| CD3<sup>+</sup>CD8<sup>+</sup> | Cytotoxic cells |
| CD3<sup>+</sup>CD8<sup>+</sup>GZB<sup>+</sup> | Activated cytotoxic T cells |
| CD3<sup>-</sup>CD68<sup>+</sup>CD209<sup>-</sup> | Macrophages |
| CD3<sup>-</sup>CD68<sup>-</sup>CD209<sup>+</sup> | Dendritic cells |
| αSMA+CD3-panCK- | Fibroblasts |
| panCK<sup>+</sup> | Cancer cells/epithelial cells |

|  |  |
| --- | --- |
| PD1 | Checkpoint receptor |
| PD-L1 | PD1 Checkpoint ligand |
| TIM3 | Checkpoint ligand |
| HLA-ABC | MHC class I |
| HLA-DR | MHC class II |
| CD45RO | Memory cells |
| IDO | Immune suppression, tryptophan catabolic enzyme, |
| Arginase | Immune suppression, Arginine catabolic enzyme |

**Table II. A. Antibody panel; B. Cell types assignments; C. Activation and other markers.**

